# Activation of 5-HT_7_ receptors in the mouse dentate gyrus does not affect theta-burst-induced plasticity at the perforant path synapse

**DOI:** 10.1101/2024.09.17.613425

**Authors:** Marcin Siwiec, Bartosz Bobula, Michał Kiełbiński, Nikola Multan, Grzegorz Hess, Krzysztof Tokarski

## Abstract

**Background:** The study examined the effects of 5-HT_7_ receptor activation on GABAergic transmission within the dentate gyrus and plasticity at the glutamatergic perforant path input.

**Methods:** Immunofluorescence imaging was performed using transverse hippocampal slices from transgenic mice expressing green fluorescent protein (GFP) under the Htr7 promoter. This was followed by whole-cell patch clamp electrophysiological recordings assessing the effects of pharmacologically activating 5-HT_7_ receptors on spontaneous inhibitory postsynaptic currents recorded from dentate granule cells and hilar mossy cells — two glutamatergic neuron types present in the dentate gyrus. Extracellular recordings of field excitatory postsynaptic potentials were then performed to assess whether 5-HT_7_ receptor activation influenced theta-burst stimulation-evoked plasticity of the perforant path synaptic input.

**Results:** It was found that parvalbumin and somatostatin interneurons in the dentate gyrus expressed GFP, which suggests they express 5-HT_7_ receptors. However, activation of 5-HT_7_ receptors had no effect on GABAergic transmission targeting mossy cells or granule cells. There was also no effect of 5-HT_7_ receptor activation on perforant path plasticity either with intact or blocked GABA_A_ receptor signaling.

**Conclusion:** The presence of 5-HT_7_ receptors in a subset of parvalbumin and somatostatin interneurons in the mouse dentate gyrus could mean that they are involved in the inhibitory control of dentate gyrus activity. However, this potential effect was not evident in slice recordings of inhibitory transmission targeting principal cells and did not affect perforant path plasticity. Further experiments are needed to fully elucidate the functional role of these receptors in the dentate gyrus.

## Introduction

GABAergic transmission is a potent regulator of the plasticity of excitatory synapses [1,2]. Although the history of the research on long-term synaptic plasticity of perforant path synapses in the dentate gyrus (DG) goes back more than 50 years [3], its modulation by local DG interneurons remains incompletely understood. Perforant path synapses in the DG are capable of bidirectional plasticity (long-term potentiation, LTP, and long-term depression, LTD [4]). The primary neuronal type of the DG are small, glutamatergic granule cells forming the granular layer. In addition to several types of GABAergic interneurons [5] located mainly along the base of the granular layer and in the hilus, the DG also contains larger and less numerous glutamatergic mossy cells, which reside in the hilar region [6]. All these different neuronal types form a complex network of synaptic connections within the DG that functions as a gate for further information flow into the hippocampus proper [7,8].

An important factor involved in control and regulation of multiple functions of the hippocampal formation, including learning and memory, is serotonin (5-HT) [reviewed in 9]. Serotonergic innervation of the hippocampal formation originates mainly from the median raphe nucleus [reviewed in: 9]. It has been shown that 5-HT, acting via 5-HT_1A_ [9], 5-HT_2C_ [10], and 5-HT_4_ [11] receptors, modulates LTP in the DG. The 5-HT_7_ receptor (5-HT_7_R), the last member of the 5-HT receptor family to be identified [12–14], is abundant in the DG as well [15] but its association with particular neuronal types in the DG is unknown. 5-HT_7_Rs are significant modulators of neuronal activity. Studies in hippocampal CA1 pyramidal neurons have demonstrated that activation of 5-HT_7_Rs enhances neuronal membrane excitability and decreases spiking latency of CA1 pyramidal cells [16,17], factors that affect the release of the neurotransmitter. It is conceivable that 5-HT_7_R activation might alter the conditions for the induction of synaptic plasticity in the DG, but the possibility of an involvement of 5-HT7Rs in the induction of LTP in the DG has not yet been investigated.

Thus, to examine the pattern of 5-HT_7_R expression in the DG, in the present study, we used a transgenic mouse line expressing green fluorescent protein (GFP) under the Htr7 promoter. Following up on immunofluorescence results showing that GFP related to 5-HT_7_R expression is present predominantly in DG inhibitory interneurons, further experiments focused on elucidating the functional consequences of 5-HT_7_R activation on GABAergic control of the two excitatory cell types of the DG — granule cells and mossy cells. The final set of experiments aimed to characterize the potential effects of 5-HT_7_R activation on the plasticity of the medial perforant path input to the DG.

## Materials and Methods

### Animals

The experimental protocols were approved by the 2nd Local Institutional Animal Care and Use Committee at the Maj Institute of Pharmacology, Polish Academy of Sciences in Krakow (document ID: 44/2020). All experiments were carried out in accordance with the European Communities Council Directive of September 22, 2010 (2010/63/UE) on the protection of animals used for scientific purposes and national law. Male wild-type and transgenic C57BL/6J mice (8-12 weeks old) were maintained on a 12/12 h light/dark schedule with standard food and tap water available *ad libitum*. The 5-HT_7_-GFP strain used in the study — Tg(Htr7-EGFP)ST29Gsat/Mmucd — was generated as part of the GENSAT project [18].

### Tissue preparation, immunofluorescent staining, and imaging

Male 5-HT_7_R-GFP mice were anesthetized using 100 mg/kg sodium pentobarbital (Biowet) and transcardially perfused with PBS followed by 4% paraformaldehyde (PFA) in 0.1M phosphate-buffered saline (PBS). After a post-fixation period of 12h at 4 °C, 50 μm-thick transverse brain sections were cut using a Leica VT-1200S vibrating blade microtome (Leica Microsystems). Free-floating sections were incubated in a blocking buffer containing 5% normal donkey serum (NDS, JacksonImmunoResearch) and 0.1% Triton X-100 (TX-100, Invitrogen) in PBS for 1h. Next, they were incubated overnight at 4°C with the following primary antibodies diluted in the blocking buffer: 1:1000 chicken anti-GFP (Aves Labs GFP-1020), 1:200 mouse anti-somatostatin (GeneTex SOM-018), 1:1000 rabbit anti-parvalbumin (PV27, SWANT). Another combination of primary antibodies used was 1:1000 chicken anti-GFP with 1:1000 mouse anti-calretinin (6B3, SWANT) and 1:1000 rabbit anti-pro-CCK (107-115aa, Frontier Institute). After extensive 3 × 10 min washing in PBS, the slices were further incubated in a mixture of the following secondary antibodies for 1h at room temperature: goat anti-chicken Alexa Fluor Plus 488, goat anti-mouse Alexa Fluor Plus 555, goat anti-rabbit Alexa Fluor Plus 647 (all from Invitrogen). Subsequently, sections were washed in PBS, counterstained with DAPI, washed again, mounted, and coverslipped with Prolong Glass antifade medium (Invitrogen). Optical z-stacks were then captured with a 20x/0.8 air objective (Plan-Apochromat 20x/0.8 M27) and Axiocam 506 camera on a Zeiss Axio Imager Z2 fluorescence microscope equipped with the Apotome 2 optical sectioning module (Carl Zeiss, Germany).

The proportion of GFP-expressing parvalbumin (PV^+^) and somatostatin (SOM^+^) interneurons was assessed using microphotographs from 3 mice (5-6 sections per animal). Images were automatically processed with FIJI/ImageJ scripts [19]. The workflow consisted of per-channel background subtraction (rolling ball method), followed by automated local thresholding. Bernsen, IsoData and moments segmentation algorithms were used for PV, SOM, and GFP signals, respectively, based on empirical testing in example images. Thresholded images were size-filtered to retain objects corresponding to intact and fully stained neuronal somata. Colocalization was assessed separately in each optical slice using the pixel value product method, followed by manual verification and, finally, z-projection and cell counting.

### Brain slice preparation for electrophysiological experiments

Mice were decapitated under isoflurane anesthesia (Aerrane, Baxter). Their brains were quickly removed and placed in cold neuroprotective artificial cerebrospinal fluid (NMDG-HEPES ACSF) containing (in mM): 92 NMDG, 2.5 KCl, 0.5 CaCl_2_, 10 MgSO_4_, 1.2 NaH_2_PO_4_, 30 NaHCO_3_ 20 HEPES, 25 glucose, 5 sodium ascorbate, 2 thiourea, 3 sodium pyruvate, pH titrated to 7.3 with HCl. Transverse 350 μm slices containing the ventral/intermediate dentate gyrus were cut on a vibrating microtome (Leica VT1200S, Leica Microsystems) and incubated at 35°C for 25 min in NMDG-HEPES aCSF while gradually introducing NaCl [20]. Afterward, slices were transferred to the HEPES holding ACSF containing (in mM): 92 NaCl, 2.5 KCl, 1.2 NaH_2_PO_4_, 30 NaHCO_3_, 20 HEPES, 25 glucose, 5 sodium ascorbate, 2 thiourea, 3 sodium pyruvate, 2 MgSO_4_, 2 CaCl_2_, pH titrated to 7.3 with NaOH, and held at room temperature for at least one hour before recording. ACSF was continuously bubbled with 95% O_2_ and 5% CO_2_. Solution osmolality was 300-310 mOsm. All compounds for ASCF solutions were purchased from Sigma-Aldrich.

### Whole-cell patch clamp recordings of spontaneous inhibitory postsynaptic currents

Slices were placed in the recording chamber and superfused at 4-5 ml/min with warm (32 ± 0.5°C) recording ACSF composed of (in mM): 132 NaCl, 2 KCl, 2.5 CaCl_2_, 1.3 MgSO_4_, 1.2 NaH_2_PO_4_, 24 NaHCO_3_ and 10 glucose, continuously bubbled with a mixture of 95% O_2_ and 5% CO_2_. Neurons were visualized on a Nikon Eclipse Fn1 upright microscope (Nikon Europe B.V.) equipped with 900 nm IR DIC optics, a 40×0.8 NA water immersion objective, and sCMOS camera (CellCam Kikker, Cairn Research), along with a computer-controlled piezo xy stage and micromanipulators (Sensapex). Patch pipettes were pulled from 1.5 mm OD/0.86 mm ID borosilicate glass capillaries (Sutter Instruments) using a P87 puller (Sutter Instruments). Spontaneous inhibitory postsynaptic currents (sIPSCs) were recorded from dentate granule cells as well as hilar mossy cells in whole-cell voltage clamp mode using a Multiclamp 700B amplifier (Molecular Devices). Signals were low-pass filtered at 2 kHz and digitized at 20 kHz with a National Instruments PCIe-6353 data acquisition card controlled by the open-source ACQ4 software package [21]. Biocytin (0.3%, HelloBio) was included in the recording pipette solution for later verification of cell morphology. A high-chloride recording pipette solution was used for recording sIPSCs and contained (in mM): 130 KCl, 5 NaCl, 0.3 CaCl_2_, 2 MgCl_2_, 10 HEPES, 10 phosphocreatine-Na_2_, 5 Na_2_-ATP, 0.4 Na-GTP and 0.3 EGTA. Osmolarity and pH were adjusted to 290 mOsm and 7.2 using sucrose and KOH respectively. Recording pipettes had open tip resistances of approximately 4-6 MΩ when filled with this solution. The liquid junction potential was experimentally verified to be -0.56 mV [22]. The recording protocols were not corrected for this minimal offset. The AMPA/kainate receptor blocker NBQX (10 μM, HelloBio) was included in the recording ACSF to isolate GABAergic synaptic activity. A holding potential of -76 mV was used to reliably record sIPSCs with few/no action potential currents present. Cell identification was based on anatomical location (mossy cells — dentate gyrus hilar region; granule cells — the outer third part of the granule cell layer in the supra-pyramidal blade of the ventral/intermediate dentate gyrus), electrophysiological properties, and cell morphology. Typical dentate granule cells had a small ovoid cell body and were characterized by a relatively hyperpolarized resting membrane potential recorded right after break-in (typically ∼ −80 mV). They also displayed a typical membrane resistance of 200-300 MΩ and low intrinsic excitability (Fig. 2A-C) [23]. Hilar mossy cells, on the other hand, typically had large, triangular, or trapezoidal cell bodies. Upon break-in, they were characterized by a large cell capacitance, membrane resistance of 150-300 MΩ, relatively higher excitability and high magnitude, frequent spontaneous synaptic activity (Fig. 3A-C) [24]. Access resistance had to stay below 20 MΩ and not change by more than 20%, and holding current had to remain stable for the entire baseline period, otherwise the recording was discarded. After recording cells were further confirmed to belong to these two neuronal populations based on dendritic morphology revealed by biocytin labeling.

### Morphological verification of recorded cells

Slices were fixed in 4% PFA in PBS for 1 h. After several washing steps, they were incubated in PBS containing 0.1% TX-100 and Alexa Fluor 647-conjugated streptavidin (1:500, Invitrogen) for 2 h at RT on an orbital shaker. After washing in PBS and counterstaining with DAPI (Invitrogen) slices were mounted on glass slides and coverslipped using the Vectashield mounting medium (Vector Laboratories). Verification of individual neuron morphology was performed after acquiring z-stacks taken with a 20×0.8 NA air apochromatic objective on a Zeiss Axio Imager Z2 fluorescence microscope using the Apotome 2 module (Carl Zeiss).

### Extracellular recordings of medial perforant path-dentate gyrus plasticity

Extracellular recordings were performed using the same ACSF and hardware as the patch clamp recordings. The WinLTP software package was used for the recording of plasticity experiments [25]. In a subset of recordings 100 μM picrotoxin (HelloBio) was added to the ACSF to block inhibitory transmission via GABA_A_ receptors. Signals were low-pass filtered at 2 kHz and high-pass filtered at 0.3 kHz. Field excitatory postsynaptic potentials (fEPSPs) were elicited using a concentric bipolar stimulation electrode (FHC) placed in the middle of the dentate gyrus molecular layer of the suprapyramidal blade, corresponding to the medial perforant path. The recording electrode (3-5 MΩ patch clamp recording pipette filled with recording ACSF) was placed approximately 200-400 μm away from the stimulating electrode (Fig. 4). Monosynaptic fEPSPs were elicited every 20 s with 200 μs wide rectangular current pulses using a constant-current stimulus isolation unit (WPI). After a 10-15 min stabilization period, the stimulation intensity was adjusted to evoke fEPSPs with a rising slope approximately 50% of the maximum value (typically 20-30 μA). A theta-burst stimulation (TBS) protocol was used to induce plasticity after recording a 20-minute baseline period. TBS consisted of a series of 5 high-frequency 100 Hz trains of 200 μs long square current pulses repeated 5 times every 200 ms. This was repeated 5 times every 10 s. The TBS protocol was chosen for a physiologically plausible LTP induction mechanism [26]. After TBS the recordings continued for 1 h with baseline stimulation intensity.

### Pharmacological activation of 5-HT_7_ receptors

Drugs were bath-applied in final concentrations for both patch clamp and extracellular recordings. The selective 5-HT_7_R agonist LP-211 (Sigma-Aldrich) was dissolved in dimethyl sulfoxide (DMSO, HelloBio) and diluted in ACSF to a final concentration of 1 μM (LP-211) and 0.1% v/v (DMSO). LP-211 has been extensively used in functional studies of 5-HT_7_Rs [27–31]. Control and/or baseline recordings were performed in 0.1% DMSO alone. The highly selective 5-HT_7_R antagonist SB 269970 (2 μM, HelloBio) was used in a subset of recordings to confirm the selectivity and specificity of any LP-211-evoked effects. The slice was perfused with the antagonist for at least 15 min before commencing recordings.

In patch clamp experiments the effect of LP-211 on GABAergic transmission was evaluated relative to baseline levels in each cell. After obtaining the whole-cell configuration and verifying the electrophysiological phenotype of the recorded cell in current clamp mode using a series of hyper- and depolarizing current pulses, the recording was switched to voltage clamp at -76 mV and after several minutes of stabilization, a 4 min baseline period of sIPSC activity was recorded in ACSF containing 10 μM NBQX and 0.1% DMSO. Next, the perfusion line was switched to ACSF containing 10 μM NBQX with 1 μM LP-211 in 0.1% DMSO and the slice was perfused for 12 min to allow the compound to activate the receptors and equilibrate before recording another 4 min period. For extracellular experiments, 1 μM LP-211 was added to the ACSF and allowed to perfuse the slice approximately 1 h before the planned induction of plasticity, i.e. 30 min before starting the recording. In control recordings 0.1% DMSO alone was used within the same time frame.

### Electrophysiology data analysis and statistics

For patch clamp experiments and sIPSC detection, the 4 min “baseline” and “LP-211” segments were analyzed to detect sIPSCs using a deconvolution algorithm [32] implemented in the Eventer software package [33]. Briefly, after creating an appropriate template separately for each cell based on representative sIPSCs and running the FFT-based deconvolution algorithm, the event cut-off was set to > 3.4 standard deviations from the histogram of the deconvoluted wave, based on experimenter judgment of false-positive/false negative rates. The resulting individually detected sIPSCs were screened for obvious false positives after an amplitude threshold cut-off at 5 pA. Data sets containing voltage clamp sIPSC frequencies and amplitudes were analyzed using an effect size estimation framework along with a bootstrap-based inference approach, as described in [34]. That is, in order to control for sample size and sampling-induced deviations from normality, 5000 bootstrap estimates of sample mean differences were derived. Next, the 95% confidence intervals were constructed from the resulting bootstrapped distributions and then were further bias-corrected and accelerated. Assessing the significance of the difference was then performed by verifying whether the 95% confidence interval of the distribution of mean differences included the zero value. If not, then the difference was deemed statistically significant. To satisfy a common requirement of scientific journals, a permutation t-test of effect size was also performed, where the p-value reported is the probability of observing the effect size (or greater), assuming the null hypothesis of zero difference is true. For each permutation p-value, 5000 reshuffles of the control and test labels were performed. We have used the open source R package *dabestr* to perform the necessary calculations [34]. This statistical approach was shown to result in improved robustness of inference to sampling variability and is much more stringent in rejecting false-positives.

Extracellular recordings were analyzed using WinLTP software [25], where three consecutive fEPSP traces (stimulation every 20 s) were averaged to obtain one low-noise reading of the fEPSP slope for every minute of the recording. The maximum slope was calculated from the rising slope of the fEPSP in a sliding 1.5 ms time window. The data were then normalized to the mean slope of the 20-minute baseline period for each recording separately. The final values for plasticity magnitude were thus expressed as percentages of the mean baseline slope level. Statistical data analysis and graphing of the results were performed using the R programming language and RStudio IDE [35,36]. R packages used were the tidyverse collection [37] and patchwork [38]. Statistics on sIPSC frequency and amplitude before and after LP-211 administration were performed using a two-tailed paired-samples t-test. Due to the bi-directionality of plasticity results and resulting lack of normality, extracellular recordings were analyzed by comparing whole distributions of plasticity magnitudes between the control and experimental groups using the nonparametric two-sample Kolmogorov-Smirnov test. The resulting distributions were plotted as cumulative probability histograms using the ecdf() R function. For all statistical procedures, the inference was two-tailed and performed with the arbitrary significance threshold set at p < 0.05. Extracellular summary fEPSP data were plotted as mean ± SEM.

## Results

### Htr7 promoter-dependent GFP is expressed in parvalbumin and somatostatin interneurons in the mouse dentate gyrus

The observed GFP immunostaining pattern in 5-HT_7_R-GFP mice consisted of sparse, but distinct, labeling of neuronal cell bodies, which, in transverse sections of the dentate gyrus, were primarily located in the hilus, with only single immunopositive cells present in the granule and molecular layers. These GFP^+^ cells were presumptive inhibitory interneurons predominantly co-expressing SOM or PV (Fig. 1), but typically not calretinin or pro-cholecystokinin (supplementary Fig. 5 and Fig. 6). For the latter two markers, only a handful of cells with weak fluorescent signal corresponding to putative GFP expression were found (not shown). Quantification of PV and SOM cell bodies revealed GFP co-expression was widespread in these populations, as double-immunopositive cells constituted 59% (223/379) of all PV^+^ and 70% (271/382) of all SOM^+^ neurons.

**Figure 1.**
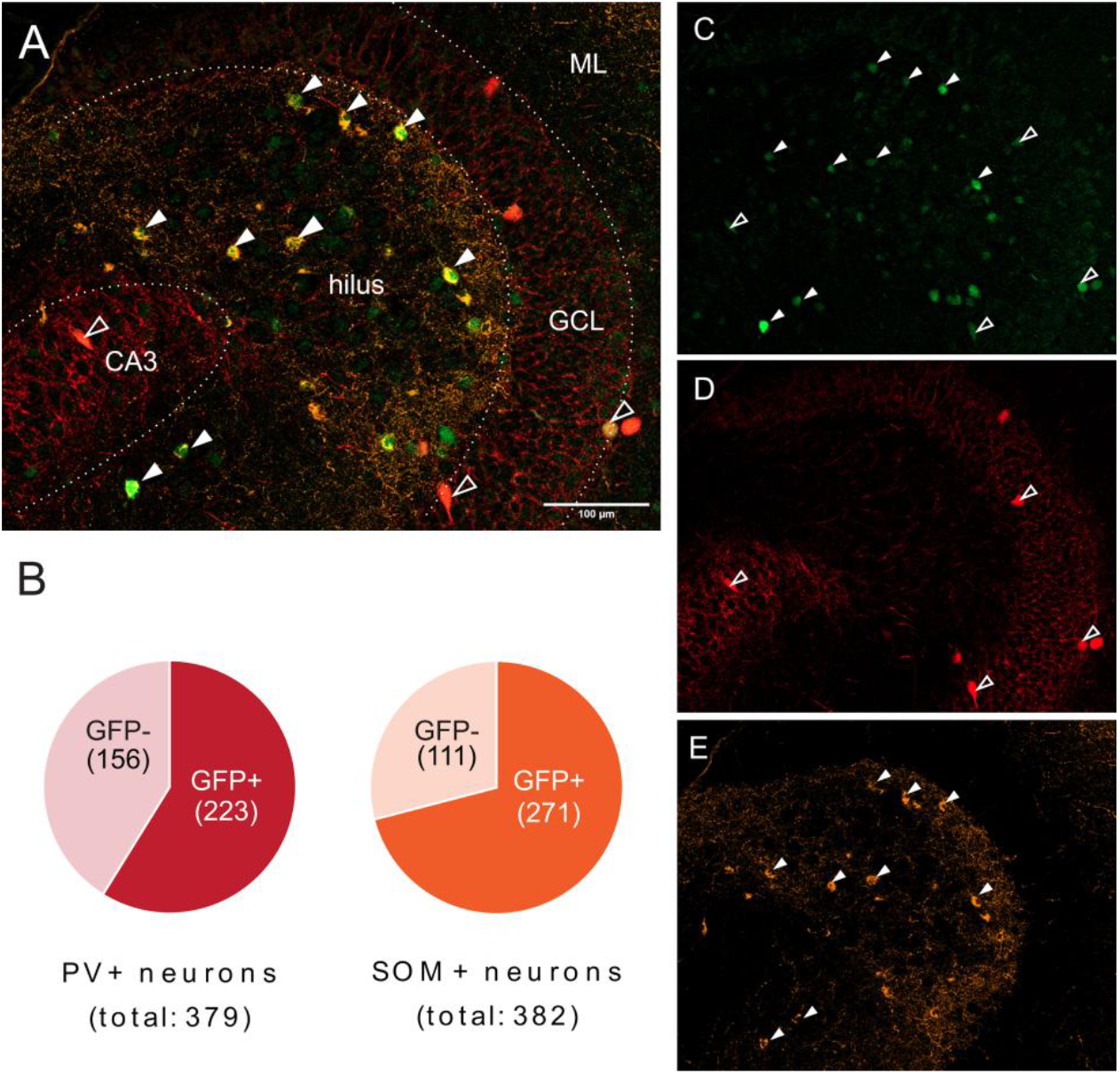
(A) Representative composite frame image of a mouse dentate gyrus section stained for GFP, SOM and PV. (B) Total counts and proportions of PV- and SOM-immunopositive cells expressing or lacking the expression of GFP across images from n = 3 mice (5-6 sections per mouse). (C-E) Single channels from image A corresponding to GFP (green; C), SOM (yellow; D) and PV (red; E). Cell bodies immunopositive for SOM–GFP and PV–GFP indicated with solid arrowheads and contour arrowheads, respectively. Abbreviations: GFP — green fluorescent protein; PV — parvalbumin; SOM — somatostatin; GCL — granule cell layer; CA3 — cornu Ammonis 3; ML — molecular layer.

### Activation of 5-HT_7_ receptors does not influence spontaneous GABAergic transmission targeting either granule cells or mossy cells

Since GFP related to 5-HT_7_R was expressed predominantly in inhibitory interneurons, further experiments focused on functional consequences of 5-HT_7_R activation on the GABAergic input to the two DG excitatory cell types. Whole-cell recordings of sIPSCs in granule cells revealed that activation of 5-HT_7_Rs by LP-211 did not influence the inhibitory transmission targeting this cell type (Fig. 2). LP-211 did not affect sIPSC frequency (Fig. 2D), as the paired mean difference in frequency against baseline for repeated measures between baseline and LP-211 was 0.77 Hz [95% CI: −2.54, 4.69; n = 31; p = 0.9332, two-sided permutation t-test]. There were no changes in amplitude either (Fig. 2E), as revealed by the paired mean difference in amplitude for repeated measures against baseline between baseline and LP-211 of 1.47 Hz [95% CI: −1.26, 4.29; n = 31; p = 0.2681, two-sided permutation t-test].

**Figure 2.**
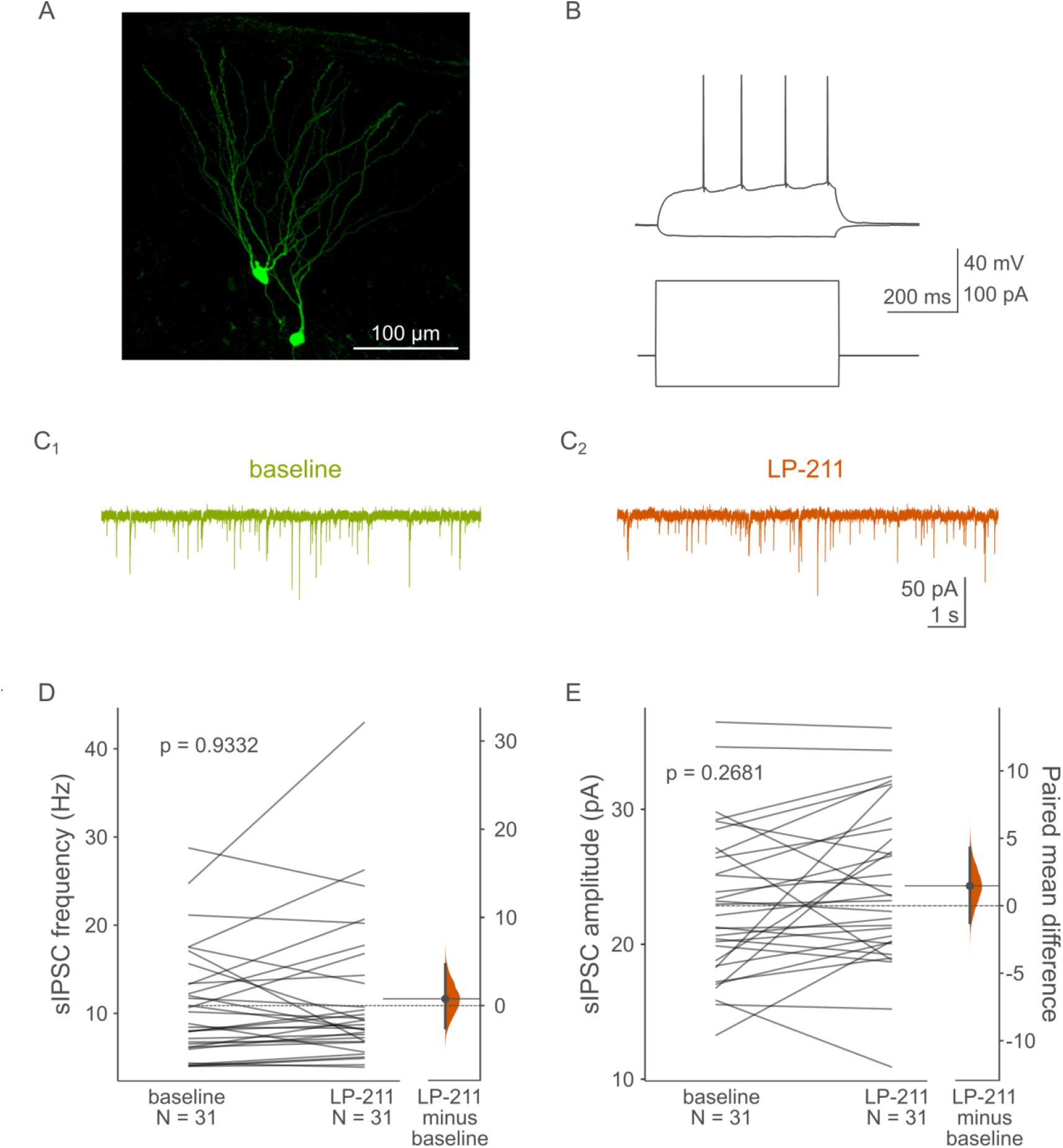
Effects of LP-211 on GABAergic transmission targeting dentate gyrus granule cells. (A) Maximum projection image of representative biocytin-filled granule cells. (B) Raw current clamp traces (top) and corresponding current steps (bottom) showing a voltage response of a typical dentate granule cell to hyperpolarizing and suprathreshold depolarizing current steps. (C1) Raw voltage clamp sIPSC traces before and after (C2) LP-211 administration. (D) Application of 1 μM LP-211 did not affect sIPSC frequency (p = 0.9332, paired two-sided permutation t-test) or (E) amplitude (p = 0.2681, paired two-sided permutation t-test). Abbreviations: sIPSC — spontaneous inhibitory postsynaptic current.

Recordings from mossy cells also did not reveal any LP-211-induced changes in sIPSC frequency (Fig. 3D) — the paired mean difference in frequency against baseline for repeated measures between baseline and LP-211 was 5.65 Hz [95% CI: −1.03, 15.57; n = 21; p = 0.272, two-sided permutation t-test]. There was also no meaningful paired mean difference in amplitude, which in this case was −0.12 Hz [95% CI: −10.48, 8.35; n = 21; p = 0.842, two-sided permutation t-test] (Fig. 3E).

**Figure 3.**
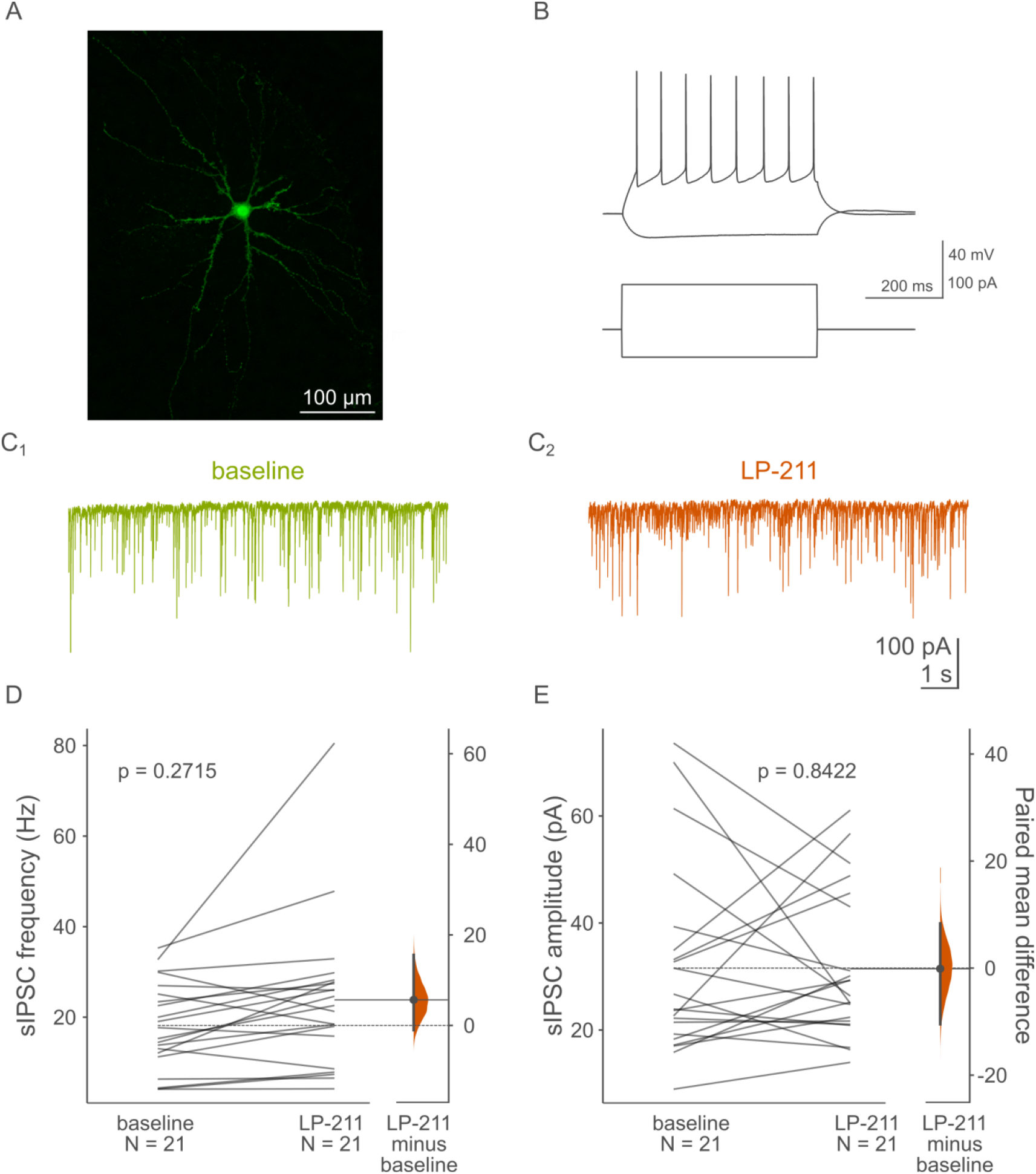
Effects of LP-211 on GABAergic transmission targeting hilar mossy cells. (A) Maximum projection image of a representative biocytin-filled hilar mossy cell. (B) Raw current clamp traces (top) and corresponding current steps (bottom) showing a voltage response of a typical hilar mossy cell to hyperpolarizing and suprathreshold depolarizing current steps. (C1) Raw voltage clamp sIPSC traces before and after (C2) LP-211 administration. (D) Application of 1 μM LP-211 did not affect sIPSC frequency (p = 0.2715, paired two-sided permutation t-test) or (E) amplitude (p = 0.8422, paired two-sided permutation t-test). Abbreviations: sIPSC — spontaneous inhibitory postsynaptic current.

### Activation of 5-HT_7_ receptors does not affect the theta-burst-evoked plasticity of the medial perforant path-granule cell synapse

To characterize the potential effects of 5-HT_7_R activation on the plasticity of the medial perforant path input to the DG, fEPSPs evoked by stimulation of the medial perforant path were recorded (Fig. 4A). In conditions with intact GABAergic signaling TBS failed to induce LTP with the majority of recordings resulting in no LTP or even LTD and the mean plasticity magnitude of 85.3 ± 8.32% relative to baseline (i.e. 100%; Fig. 4 B1-B2). Activation of 5-HT_7_Rs by preincubating slices with ACSF containing 1 μM LP-211 did not change this distribution of plasticity magnitudes (N_control_ = 17, N_LP-211_ = 17; D_17,17_ = 0.23529, p = 0.75061, exact two-sample Kolmogorov-Smirnov test), with the mean plasticity magnitude in the LP-211 group of 87.5 ± 5.13%. Exactly the same plasticity induction protocol resulted in a robust LTP when 100 μM picrotoxin (PTX) was added to the ACSF to block fast GABAergic inhibition (Fig. 4 C1-C2), with the mean plasticity magnitude of 153 ± 24%. This time the distribution of plasticity magnitudes strongly favored LTP over LTD, however, when LP-211 was included in the recording ACSF the distribution of plasticity magnitudes remained unchanged (N_control_ = 19, N_LP-211_ = 13; D_19,13_ = 0.27126, p = 0.51867, exact two-sample Kolmogorov-Smirnov test), with the mean plasticity magnitude of 166 ± 28% (Fig. 4 C1-C2.)

## Discussion

Hippocampal neurons express a multitude of serotonin receptors and their associated molecular effectors. To date, many serotonin receptor subtypes have been identified in the dentate gyrus including, but not limited to 5-HT_1A_, 5-HT_1B_, 5-HT_1D_, 5-HT_2A_, 5-HT_2C_, 5-HT_3_, 5-HT_4_, 5-HT_5A_, 5-HT_6_ and 5-HT_7_ [11,39–47]. For some of these receptors, such as the 5-HT_7_R, the contribution to overall hippocampal physiology is yet to be defined. The results of this study indicate that in the dentate gyrus 5-HT_7_Rs are expressed in inhibitory interneurons of the hilar region with GFP present in the majority of PV^+^ and SOM^+^ neurons in the area. However, activation of these receptors did not enhance the inhibitory synaptic input to either mossy cells or granule cells.

It has been shown that mossy cells function as local circuit integrators and exert modulatory influence on dentate granule cells as well as the CA3 through back-projecting pathways [48,49]. They receive considerable excitatory synaptic input from granule cells’ massive specialized synaptic boutons (mossy terminals) apposed to thorny excrescences present on their cell bodies and proximal dendrites, giving them their “mossy” appearance [50]. In turn, mossy cells are monosynaptically coupled to granule cells both ipsi- and contralaterally. However, this connection is only weakly excitatory, prone to synaptic failures and strongly masked by GABAergic inhibition. Therefore, it is believed that the primary effect of mossy cell activation on granule cells is in fact non-direct and inhibitory, via the excitation of local GABAergic interneurons [51]. Some studies suggest that this interaction can become reversed in pathological states of hippocampal circuit activity such as those resulting from acute trauma and epileptogenesis [52,53]. Furthermore, direct excitatory connections have also been discovered between individual mossy cells, which could in theory further consolidate their influence on granule cell activity [54].

Given the considerable glutamatergic drive via mossy terminals, the activity of mossy cells has to be tightly controlled by various subtypes of dentate and hilar interneurons [55–57]. It is likely that GABAergic input to mossy cells is at least partially provided by hilar somatostatin-positive neurons [50,58]. Activation of 5-HT_7_Rs by LP-211 would, therefore, at least in principle, be expected to increase GABAergic transmission targeting mossy cells. This would probably be a direct result of increased spiking activity of these interneurons, as 5-HT_7_Rs are known enhancers of neuronal excitability [16,17,59,60]. To date, there is limited data about the interconnectivity between hilar somatostatin interneurons and mossy cells. A recent investigation of somatostatin interneuron projections to the contralateral hippocampus showed that traced somatostatin axons form some varicosities in close proximity to mossy cells, among other hilar cell types. However, they do so at very low frequencies, with these types of connections being about 24-fold less common compared to those formed by hilar mossy cells [61]. Local connections, however, should be more common, especially when considering that a subpopulation of hilar somatostatin interneurons has extensive axonal branching in the hilar region [58]. Somatostatin interneurons are also known regulators of granule cell activity and therefore the question remains as to why we did not record increased GABAergic transmission in either one of the dentate glutamatergic cell types.

One possible explanation is that the majority of dentate gyrus somatostatin interneurons are the so-called hilar perforant path-associated (HIPP) interneurons. They are characterized by weak, slow and unreliable inhibition of granule cell distal dendrites in the outer molecular layer. This is in contrast to the strong, fast and reliable perisomatic inhibition provided by parvalbumin basket cells [57]. Any increase in somatostatin interneuron spiking activity would have to be strong enough to overcome the weak synaptic coupling in order to be reliably detected with somatic granule cell recordings. Recently, however, a second subpopulation of dentate somatostatin interneurons was characterized in addition to HIPP cells. These so-called hilar interneurons (HILs) extensively innervate the dentate hilar region where mossy cell somata are located [58]. Further research is needed to characterize the degree of connectivity between these two neuron types, which could help answer why putative activation of 5-HT_7_Rs on HIPP/HIL cells does not result in increased GABAergic tone recorded from mossy cells.

There are limited sources in the literature concerning if and how parvalbumin-positive interneurons innervate hilar mossy cells. However, the majority of parvalbumin interneurons in the dentate gyrus are basket cells characterized by large triangular cell bodies in the subgranular layer as well as extensive, basket-like branching of axon collaterals around granule cell somata. They have few axons present in the hilar region [57]. As for the rest of the parvalbumin interneuron cell population, not much is known about their postsynaptic targets and these may in fact include mossy cells. However, it is important to note that in general, hippocampal parvalbumin interneurons have different electrophysiological characteristics compared to their somatostatin counterparts. Although they are usually fast-spiking neurons, they have a lower input resistance and a more hyperpolarized resting membrane potential [57]. They are often quiescent under standard *ex vivo* slice recording conditions. Therefore activation of 5-HT_7_Rs located on these cells might not provide a depolarization strong enough for spontaneous spiking to occur, at least not in standard, baseline conditions such as those employed in this *ex vivo* study. Given that we demonstrate robust 5-HT_7_R-GFP labeling of dentate parvalbumin interneurons, future experiments would need to study the possible functional implications of 5-HT_7_R activation *in vivo* or during *in vivo*-like activity patterns, evoked pharmacologically or electrically, where different classes of GABAergic interneurons would be driven to spiking via innate circuit activity.

One such attempt to recapitulate such mechanisms *ex vivo* was made in this study, where we performed extracellular recordings measuring the synaptic plasticity of the medial perforant path input to the dentate gyrus. A theta-burst stimulation protocol was chosen (Fig. 4), as it reliably evokes hippocampal LTP and is physiological in nature, as it attempts to mimic theta frequency hippocampal oscillations, in contrast to the more commonly used high-frequency stimulation paradigm [26,62,63]. The dentate gyrus perforant path input is under strong GABAergic control and without blocking GABA_A_ receptors it is highly resistant to LTP and even prone to LTD, as shown in our experiments (Fig. 4B1-B2). However, it was in principle possible for the 5-HT_7_R-evoked increase in GABAergic inhibition of mossy cells to shift the distribution of plasticity magnitudes, which was what we expected to occur. Somewhat surprisingly, preincubation of slices in ACSF containing LP-211 did not affect the distribution of recorded plasticity magnitudes with intact as well as blocked fast GABAergic transmission. One possible explanation for this lack of effect is that, as we demonstrated, there were no changes in GABAergic transmission recorded in mossy and granule cells following 5-HT_7_R activation. Mossy cells are potent regulators of granule cell activity. However, they usually form translaminar or even contralateral connections, which are inevitably severed in transverse slice preparations typically employed to study hippocampal plasticity [50]. Another possibility is that 5-HT_7_R-expressing interneurons are simply not recruited strongly enough by the TBS protocol for the 5-HT_7_R activation to be able to reliably influence plasticity. Indeed, GABAergic transmission targeting granule cells did not change in response to 5-HT_7_R activation, and granule cell dendrites are the main source of the fEPSP signal in extracellular recordings of perforant path plasticity.

**Figure 4.**
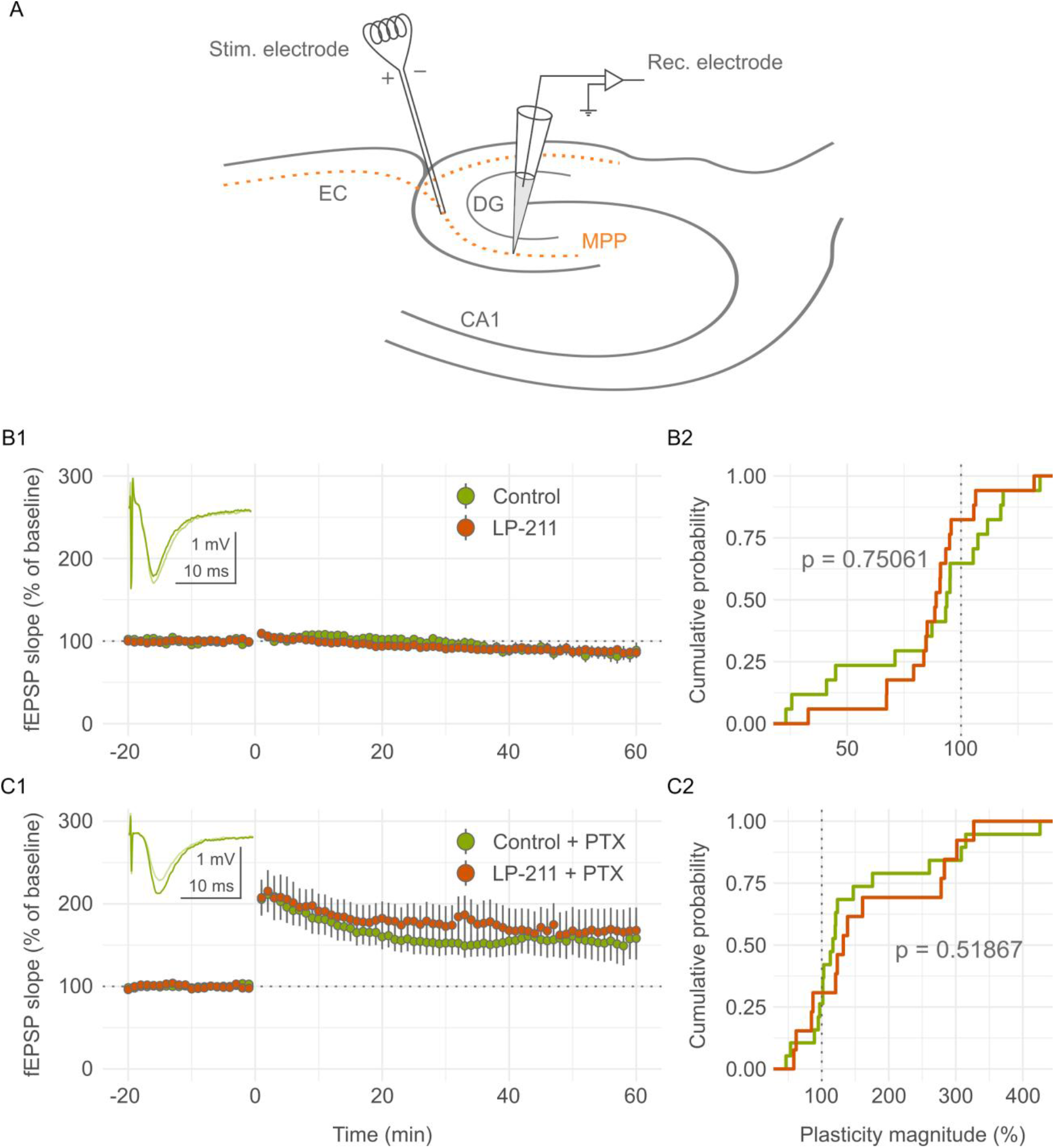
Effects of LP-211 on the magnitude of medial perforant path-dentate gyrus plasticity. (A) Anatomical diagram showing the placement of recording and stimulation electrodes. Abbreviations: EC — entorhinal cortex; CA1 — cornu Ammonis 1; DG — dentate gyrus; MPP — medial perforant path. (B1) Effects of LP-211 preincubation on fEPSP slope (%) before and after TBS. Inset: raw fEPSP traces of a representative control recording before (lighter green) and after (darker green) TBS. (B2) Cumulative probability histograms showing that preincubation of slices with LP-211 did not affect the distribution of plasticity magnitudes (N_control_ = 17, N_LP-211_ = 17; p = 0.75061, exact two-sample Kolmogorov-Smirnov test). (C1) Effects of LP-211 preincubation on fEPSP slope (%) before and after TBS with 100 μM picrotoxin (PTX) present in the ACSF to block GABAA receptors. Inset: raw fEPSP traces of a representative control recording before (lighter green) and after (darker green) TBS. (C2) Cumulative probability histograms showing that preincubation of slices with LP-211 with PTX present in the ACSF did not affect the distribution of plasticity magnitudes (N_control_ = 19, N_LP-211_ = 13; p = 0.51867, exact two-sample Kolmogorov-Smirnov test). Abbreviations: fEPSP — field excitatory postsynaptic potential; TBS — theta-burst stimulation.

**Figure 5.**
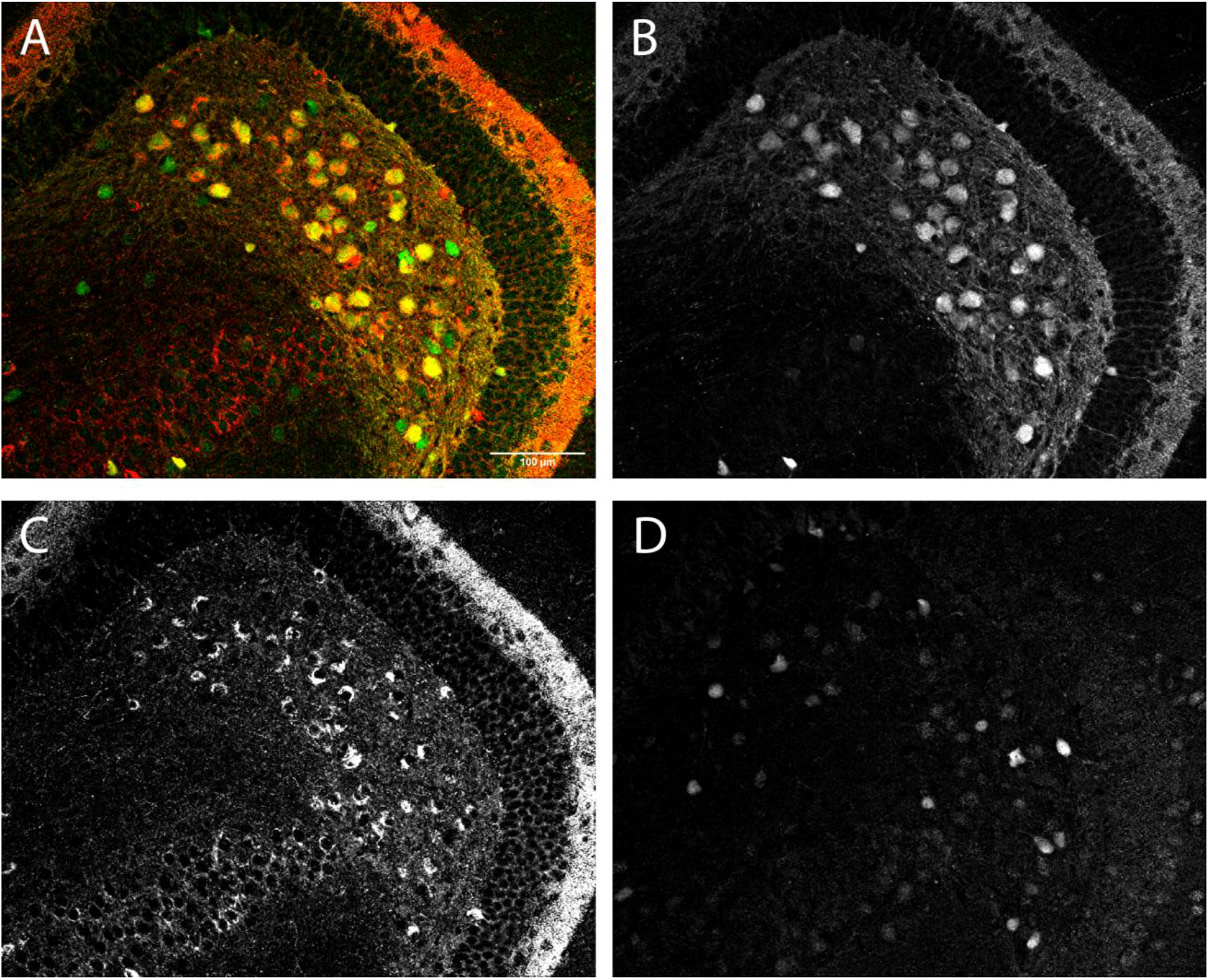
(A) Representative composite frame image of a mouse dentate gyrus section stained for calretinin (CR), pro-cholecystokinin (pCCK) and GFP. (B-D) Single channels from image A corresponding to CR (panel B), pCCK (C) and GFP (D).

**Figure 6.**
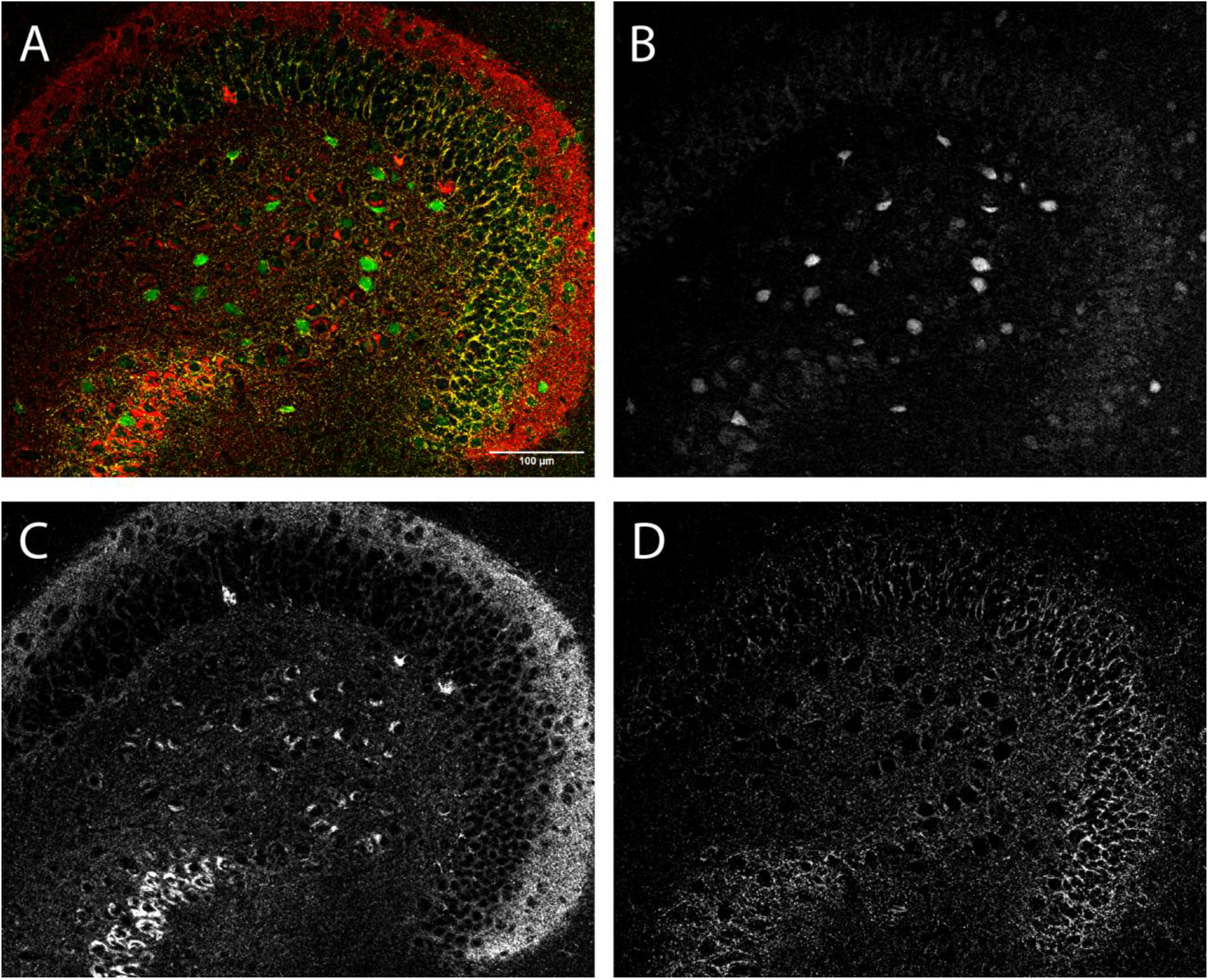
(A) Representative composite frame image of a mouse dentate gyrus section stained for GFP, pCCK and glutamate decarboxylase (GAD67). (B-D) Single channels from image A corresponding to GFP (panel B), pCCK (C) and GAD67 (D).

In conclusion, assuming that the 5-HT_7_R is at all involved in regulating dentate circuitry, it likely does so via cellular and network mechanisms not directly connected to regulating resting state GABAergic tone. Or at least not in a way that can be interrogated in standard *ex vivo* hippocampal slice preparation. Thus, this study, while providing preliminary insights into the function of 5-HT_7_Rs in the dentate gyrus, has in fact resulted in more questions than answers. For example about the functional consequences of 5-HT_7_R activation on dentate gyrus circuit activity and synaptic plasticity *in vivo*. Or the precise somatostatin and parvalbumin interneuron subtypes that express these receptors. Successfully answering these and other questions would probably require the combination of the controlled environment of *ex vivo* approaches, with pharmacological or optogenetic manipulations aimed at replicating a more natural *in vivo*-like network state.

## List of Abbreviations

5-HT: 5-hydroxytryptamine, serotonin
5-HT_7_R: 5-HT_7_ receptor
ACSF: artificial cerebrospinal fluid
AMPA: α-amino-3-hydroxy-5-methyl-4-isoxazolepropionic acid
CA1-3: cornu Ammonis 1-3
DG: dentate gyrus
DMSO: dimethyl sulfoxide
EC: entorhinal cortex
fEPSPs: field excitatory postsynaptic potentials
GABA: Gamma-aminobutyric acid
GCL: granule cell layer
GFP: green fluorescent protein
HILs: hilar interneurons
HIPP: hilar perforant path-associated
LTD: long-term depression
LTP: long-term potentiation
ML: molecular layer
MPP: medial perforant path
PBS: phosphate buffered saline
PFA: paraformaldehyde
PTX: picrotoxin PV parvalbumin
sIPSCs: spontaneous inhibitory postsynaptic currents
SOM: somatostatin
TBS: theta-burst stimulation

## Data Availability

Electrophysiology data sets generated during the current study are available from the corresponding author on reasonable request. The microscopy image dataset containing raw z-stacks has been uploaded to Zenodo and is available at https://zenodo.org/records/13919065.

## Acknowledgements

The authors would like to thank the Editor and the anonymous Reviewers for their thorough and constructive critique of this work, which, in our opinion, positively contributed to the scientific value of the manuscript.

## Author Contributions

M.S. designed the study, performed immunofluorescent staining, patch clamp, and extracellular recordings, analyzed the data, prepared Figs. 2, 3, 4, wrote and revised the manuscript. B.B. performed extracellular recordings and analyzed the data. M.K. analyzed immunofluorescence images, prepared Fig. 1, Supplementary Figs 5 and 6, helped write and revised the manuscript. N.M. performed extracellular recordings. K.T. revised the manuscript and provided guidance and supervision. G.H. helped conceptualize the study, wrote and revised the manuscript, provided guidance and supervision. All authors revised and approved the final manuscript.

## Supplementary Figures

